# IpaA reveals distinct modes of vinculin activation during *Shigella* invasion and cell-matrix adhesion

**DOI:** 10.1101/2023.03.23.533139

**Authors:** Benjamin Cocom-Chan, Hamed Khakzad, Mahamadou Konate, Daniel Isui Aguilar, Chakir Bello, Cesar Valencia-Gallardo, Yosra Zarrouk, Jacques Fattaccioli, Alain Mauviel, Delphine Javelaud, Guy Tran Van Nhieu

**Affiliations:** Team “Ca2+ signaling and Microbial Infections”, I2BC, 91190 Gif-sur-Yvette, France; Institut National de la Santé et de la Recherche Médicale U1280, 91190 Gif-sur-Yvette, France; Centre National de la Recherche Scientifique UMR9198, 91190 Gif-sur-Yvette, France; Université de Lorraine, CNRS, Inria, LORIA, F-54000 Nancy, France; Equipe Communication Intercellulaire et Infections Microbiennes, Centre de Recherche Interdisciplinaire en Biologie (CIRB), Collège de France, 75005 Paris, France; Institut National de la Santé et de la Recherche Médicale U1050, 75005 Paris, France; Centre National de la Recherche Scientifique UMR7241, 75005 Paris, France; MEMOLIFE Laboratory of excellence and Paris Science Lettre; PASTEUR, Département de Chimie, École Normale Supérieure, PSL University, Sorbonne Université, CNRS, 75005 Paris, France; Institut Pierre-Gilles de Gennes pour la Microfluidique, 75005 Paris, France; Institut Curie, PSL Research University, INSERM U1021, CNRS UMR3347, Team “TGF-ß and Oncogenesis“, Equipe Labellisée LIGUE 2016, F-91400, Orsay, France; Université Paris-Sud, F-91400, Orsay, France; Centre National de la Recherche Scientifique UMR 3347, 91400 Orsay, France

**Keywords:** vinculin, head domain, oligomerization, cell adhesion

## Abstract

Vinculin is a cytoskeletal linker strengthening cell adhesion. The *Shigella* IpaA invasion effector binds to vinculin to promote vinculin supra-activation associated with head-domain mediated oligomerization. Our study investigates the impact of mutations of vinculin D1D2 subdomains’ residues predicted to interact with IpaA VBS3. These mutations affected the rate of D1D2 trimer formation with distinct effects on monomer disappearance, consistent with structural modeling of a “closed” and “open” D1D2 conformer induced by IpaA. Notably, mutations targeting the closed D1D2 conformer significantly reduced *Shigella* invasion of host cells as opposed to mutations targeting the open D1D2 conformer and later stages of vinculin head-domain oligomerization. In contrast, all mutations affected the formation of focal adhesions (FAs), supporting the involvement of vinculin supra-activation in this process. Our findings suggest that IpaA-induced vinculin supra-activation primarily reinforces matrix adhesion in infected cells, rather than promoting bacterial invasion. Consistently, shear stress studies pointed to a key role for IpaA-induced vinculin supra-activation in accelerating and strengthening cell matrix adhesion.

## Introduction

Vinculin is a cytoskeletal linker of integrin-mediated matrix adhesions as well as cadherin-based cell-cell junctions (Goldman, 2016; Bays and De Mali, 2017). It plays an important role in cell adhesion processes, motility and development and its functional deficiency is associated with major diseases including cancer and cardiomyopathies (Peng et al., 2011). Vinculin associates with a number of ligands, including focal adhesion and intercellular junction components, lipids, signaling proteins as well as proteins regulating the organization and dynamics of the actin cytoskeleton (Goldman, 2016; Bays and De Mali, 2017). The role of vinculin in integrin-mediated adhesion has been particularly studied (Atherton et al., 2016; Parsons et al., 2010). Vinculin is recruited at focal adhesions and reinforces the link between integrins and the actin cytoskeleton. The extent of vinculin recruitment determines the growth and maturation of focal adhesions, associated with the scaffolding of adhesion components and mechanotransduction linked to the actomyosin contraction (Atherton et al., 2016; Parsons et al., 2010). While the precise mechanisms leading to vinculin activation involving combinatorial stimulation and recruitment at focal adhesions in cells are not fully understood, *in vitro* biomimetic mechanical and structure-function studies have enlightened major aspects of the role of vinculin in cell adhesion (Yan et al., 2015).

Vinculin is classically described as a three-domain protein containing an aminoterminal globular head domain (Vh), a flexible linker domain, and a F-actin-binding tail domain (Vt). Vh contains three subdomains D1-D3 and a small subdomain D4. With the exception of D4 containing a single helix bundle, each D1-D3 subdomain corresponds to a conserved repeat consisting of two four/five helix bundles connected by a long alpha-helix (Goldman, 2016). At the inactive state, vinculin is maintained folded by intramolecular interactions between Vh and Vt. All ligands activating vinculin contain a vinculin binding site (VBS), corresponding to 20-25 residues structured into an amphipathic α-helix that interacts with the first helix-bundle of D1 (Gingras et al., 2005). Insertion of the activating VBS in the D1 first helix bundle leads to the reorganization of the bundle and destabilizes the interaction between D1 and Vt (Izard et al., 2004). More than 70 VBSs have been identified in the various vinculin ligands, often containing multiple VBSs (Kluger et al., 2020). These VBSs, however, are often buried into helix bundles and their exposure regulates vinculin activation (Kluger et al., 2020). Force-induced stretching of a vinculin ligand such as talin, acting as mechanosensor, provides with a means to expose the VBSs during integrin-mediated adhesion (Sun et al., 2016; Goult et al., 2021). Talin binds to F-actin via at least two sites in its rod domain and to integrin cytoplasmic domains via its amino-terminal FERM domain. During mechanotransduction, talin stretching by the actomyosin contraction exposes vinculin binding sites (VBSs) that are buried in helix bundles of the rod domain in the native state (Yan et al., 2015). Exposed talin VBSs in turn bind to vinculin and relieve the intramolecular interactions between the vinculin head (Vh) and tail (Vt) domains, unveiling the F-actin-binding site in Vt (Goult et al., 2021). Since talin contains 11 VBSs in helix bundles unfolding at different force amplitudes, the stretching force-dependent interaction between talin and vinculin serves as a mechanism to strengthen actin cytoskeletal anchorage as a function of substrate stiffness (Yan et al., 2015; Yao et al., 2016). *In vitro*, vinculin interaction with a VBS is sufficient to promote its opening and interaction with F-actin. Vinculin activation in cells, however, may result from a combinatorial stimulus including interaction with F-actin, phosphatidylinositol (4,5)-biphosphate (PIP2) or phosphorylation (Izard and Brown, 2016; Aurnheimer et al., 2015).

Intracellular bacterial pathogens such as *Chlamydia*, *Rickettsia*, and *Shigella* express ligands diverting vinculin functions to promote virulence (Thwaites et al., 2015; Park et al., 2011; Valencia-Gallardo, 2015). Among these, the *Shigella* type III effector IpaA was shown to target vinculin via three VBSs present at its carboxyterminal domain (Valencia-Gallardo, 2015). Unlike other host cell endogenous VBSs, IpaA VBSs are not buried into helix bundles and therefore likely act in concert to promote bacterial invasion. IpaA VBS1 and VBS2 bind to the first and second bundles of D1, respectively, conferring binding to vinculin with a very high affinity and the IpaA property to act a “super-mimic” of endogenous activating VBSs (Izard et al., 2006; Tran Van Nhieu and Izard, 2007). The role of IpaA VBS3 appears more complex and likely underline the role of IpaA in different processes during bacterial invasion. In addition to vinculin, IpaA VBS3 also binds to talin and may stabilize a partially stretched talin conformer present in filopodial adhesions, thereby favoring bacterial capture by filopodia at initial stages of the bacterial invasion process (Park et al., 2011; Valencia-Gallardo et al., 2019). IpaA VBS3 was also shown to bind to the vinculin D2 subdomain, when IpaA VBSs 1-2 (aVBS1-2) are bound to D1 (Valencia-Gallardo et al., 2023). Binding of IpaA VBS1-3 (aVBD) to D1D2 triggers major conformational changes, coined “supra-activation”, leading to D1D2 homo-oligomerization via the D1D2 head subdomains and the formation of D1D2 trimers (Valencia-Gallardo et al., 2023). IpaA-induced vinculin “supra-activation” enables invasive *Shigella* to promote strong adhesion in the absence of mechanotransduction (Valencia-Gallardo et al., 2023). Additionally, analysis of a vinculin cysteine-clamp variant, deficient for vinculin supra-activation but still proficient for canonical activation, suggests that vinculin head domain oligomerization is required for vinculin-dependent actin bundling and the maturation of focal adhesions into large adhesion structures (Valencia-Gallardo et al., 2023).

Seminal studies based on rotary shadow electron microscopy analysis reported the formation of vinculin mediated by Vh-Vh as well as Vt-Vt interactions *in vitro* (Molony and Burridge, 1985). Ever since, studies on vinculin oligomerization have essentially focused on Vt-Vt interactions. Phosphatidylinositol (4, 5) bisphosphate (PIP2) binding to vinculin was shown to promote vinculin oligomerization and PIP2-binding deficient vinculin showed defects in the organization of the actin cytoskeleton, as well as increased turn-over of focal adhesions (Bakolitsa et al., 1999; Chinthalapudi et al., 2014). Binding of F-actin was also reported to induce the formation of vinculin tail dimers, likely different than those induced by PIP2, and mutants in the Vt C-terminal hairpin responsible for F-actin-induced dimerization showed defects in actin bundling associated with a decrease in size and number of focal adhesions (Johnson and Craig, 2000; Bakolitsa et al., 1999). Vinculin establishes catch bonds with a significantly increased lifetime when the force is applied towards the pointed end of actin filaments, consistent with the polarity of actomyosin contraction (Huang et al., 2017). However, while mechanotransduction triggers vinculin recruitment associated with the maturation, enlarging of focal adhesions and actin bundling, how Vt-mediated vinculin oligomerization may be regulated by actomyosin contraction remains unclear (Thompson et al., 2013). Vinculin head-domain (Vh) mediated oligomerization could provide with a force-dependent mechanism, since vinculin was reported to acts as a mechanosensor, with its head domain undergoing conformational changes under applied force, showing increased binding to ligand such as MAPK1 (Garakni et al., 2017). Here, we report the effects of mutation in vinculin D1D2 altering the formation of trimers induced by IpaA. We identified D1D2 polar residues predicted to contact IpaA VBS3 and showed their involvement in IpaA-induced trimerization. We show that these mutations affect the formation of focal adhesions providing evidence that Vh-Vh mediated vinculin oligomerization similar to that induced by *Shigella* IpaA also occurs during the maturation of cell adhesion structures.

## Results

### Design of mutations in D1D2 affecting IpaA-induced trimer formation

Structural models indicate that aVBD triggers a 30 % angle displacement of the major axis of the D1 and D2 subdomains relative to the apo D1D2 or D1D2 in complex with IpaA VBS1-2 only (Fig. 1B; Valencia-Gallardo et al., 2023). As opposed to the *closed* D1D2 conformer, this *open* D1D2 conformer is associated with D1D2-mediated trimerization of vinculin (Valencia-Gallardo et al., 2023). We hypothesized that IpaA VBS3 induced allosteric changes leading to D1D2 trimerization and that mutations in D1D2 residues interfacing IpaA VBS3 should affect trimer formation. We therefore scrutinized the interface residues between IpaA VBS3 and D1D2 in the D1D2:aVBD complex.

**Fig. 1.**
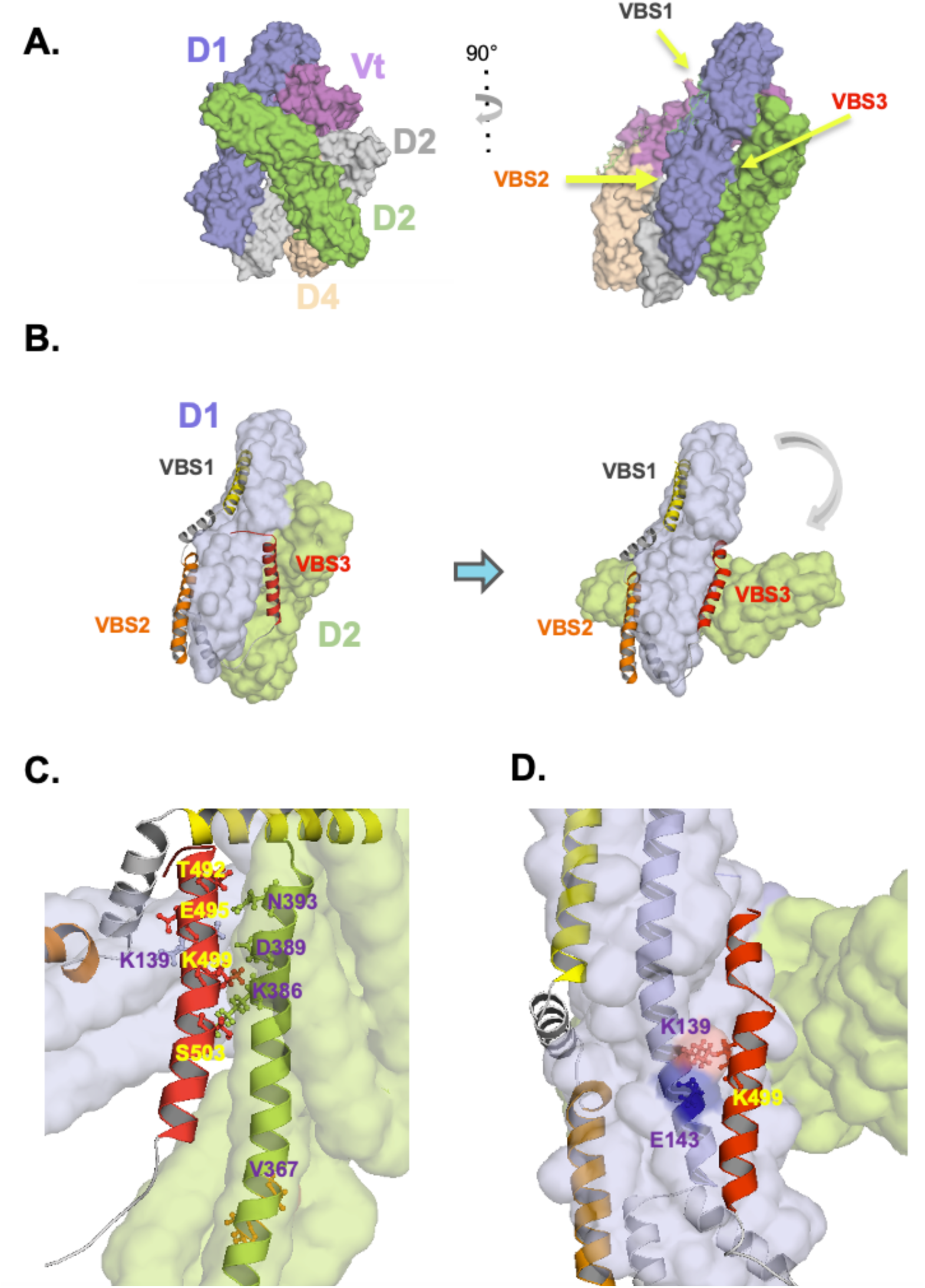
Design of D1D2 mutations targeting IpaA VBS3 contact sites. **A**, Surface structure of full-length human vinculin from the crystal structure resolved by Bakolitsa et al., 2004. The vinculin subdomains are indicated in different colors. Right: The arrows point at binding of: IpaA VBS1 to the D1 first bundle, IpaA VBS2 to the second D1 bundle, IpaA VBS3 binding to the D1D2 interface and D2 second bundle. **B-D**, cross-linking mass spectrometry-based modelling based models of D1D2: aVBD (Hauri, Khakzad et al., 2019). The vinculin D1 and D2 are shown as pale grey and pale green surface structures, respectively. IpaA VBSs are shown as ribbon structures. **B**, Left: aVBD binds to D1D2 that adopts a conformation similar to that of apoD1D2 (closed conformer). Right: binding of IpA VBS3 to the D1D2 interface induces a major conformational change (open conformer) with a 30° tilt in the relative orientation of the D1 and D2 major axis. **C, D**, higher magnifications showing the IpaA VBS3(red) interface with D1 D2 in the closed (**C**) and open (**D**) conformer. The residues potentially involved in polar interactions via their side chains are indicated. In the open conformer, IpaA K499 may interact with E143 or clash with K139 on vinculin D1.

As shown in Fig. 1B, in the closed D1D2:aVBD complex, IpaA VBS3 mainly interacts with the H10 helix of D2. A set of putative polar interactions and salt bridges can be identified, where IpaA residues T492, E495, K499 and S503 interact with vinculin residues N393, K139, D389 and K386, respectively (Fig. 1C). In the open complex, IpaA VBS3 mainly interacts with the H4 helix of D1, where IpaA K499 interacts with vinculin E143. Of note, in this open complex, an electrostatic clash between vinculin K139 and IpaA K499 may contribute to the dynamics of the IpaA VBS3 during its interaction with different allosteric conformers leading to D1D2 oligomerization (Fig. 1D). All identified vinculin residues were substituted for a charged residue to introduce a charge inversion or disrupt polar interactions using site directed mutagenesis (Materials and Methods, Table 1). In this rationale, mutations affecting IpaA VBS3’s interface with the close D1D2 conformer are expected to alter its initial docking of IpaA VBS3 on D1D2, whereas the E143K charge inversion would destabilize the open conformer and alter subsequent allosteric changes leading to trimer formation.

**Table 1.**
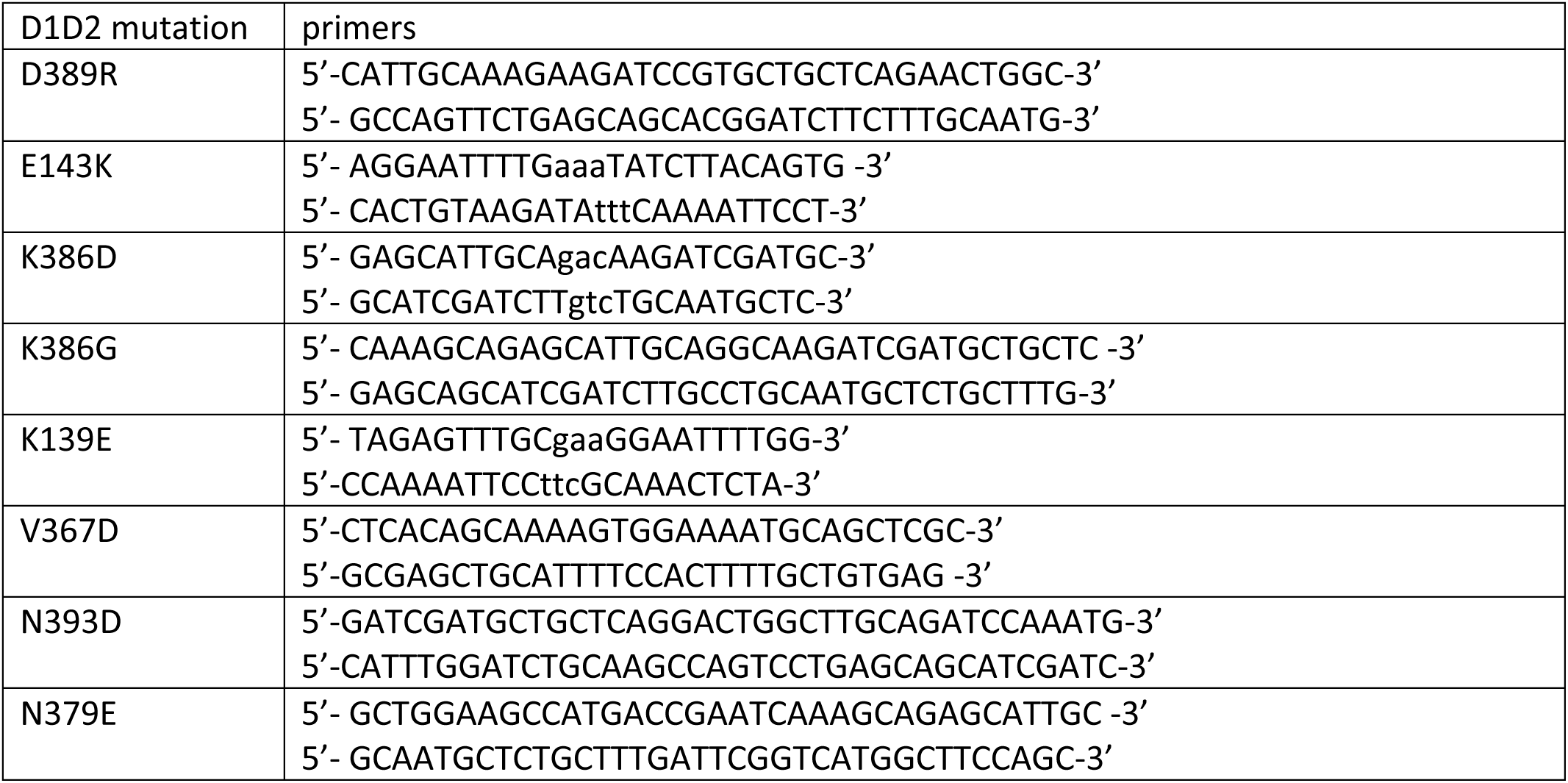
Primers used in this study.

Independent of the aVBD-D1D2 interface, the hidden Markov model-based algorithm MARCOIL predicts the presence of a coiled-coil domain in the same H10 helix of D2 between vinculin residues 348 to 393, buried into the D2 helix bundles and adjacent to the IpaA VBS3 interaction sites in the close D1D2 conformer (Delorenzi and Speed, 2002; Fig. S1). This putative coiled-coil domain contains the classical a-g heptad sequence at residues 367-373, with the V367 and A373 corresponding to the “a” and “d” hydrophobic residues, respectively, presumed to intersperse their non-polar side chains at the α-helices interfaces during oligomerization. To test the role of this domain in D1D2 trimerization, we introduced the V367D substitution predicted to disrupt supercoiled helix packing (Fig. S1).

### Quantitative analysis of D1D2 trimer formation using CN-PAGE

In previous works, we showed that aVBD induced the formation of D1D2:aVBD 1:1, as well as dimeric complexes and trimeric D1D2 complexes rapidly associating and dissociating an aVBD molecule in SEC-MALS experiments (Valencia-Gallardo et al., 2023). Here, we studied the effects of increasing aVBD molar ratio using SEC and CN (Clear Native)-PAGE. As shown in Fig. 2, at a D1D2:aVBD 1:1 molar ratio, the major peaks with similar amplitude corresponded to the 1:0 and likely the D1D2 trimeric complexes (Fig. 2A, peaks A and C). At a D1D2:aVBD 1:1.5 molar ratio, the D1D2 trimeric complexes represented the major peak (Fig. 2A, peak A). Upon increasing of IpaA molar ratio, a major shifted band (Fig. 2B, A’) could also be observed in CN-PAGE followed by Coomassie blue staining, with apo D1D2 rapidly disappearing (Fig. 2B, C’) and the formation of intermediate shifted bands (Fig. 2B). The similarity between the distribution of the bands with increasing molar ratio of aVBD in CN-PAGE and the SEC peaks suggested that the A and A’ peaks corresponded to trimeric D1D2 complexes. To confirm this, we dissected the A’ band form CN-PAGE and analyzed it in a second dimension using regular SDS-PAGE (Materials and Methods) and compared the amounts of D1D2 and aVBD relative to those present in fractions from the SEC peak A. As shown in Fig. 2C, the relative amounts if D1D2 and AVBD were similar for both A and A’ samples, with an estimated molar ratio of 3.8 and 3.2, respectively, both values being consistent with a mixture of D1D2:aVBD 3:0 and 3:1 complexes inferred from the SEC-MALS analysis (Valencia-Gallardo et al., 2023).

**Fig. 2.**
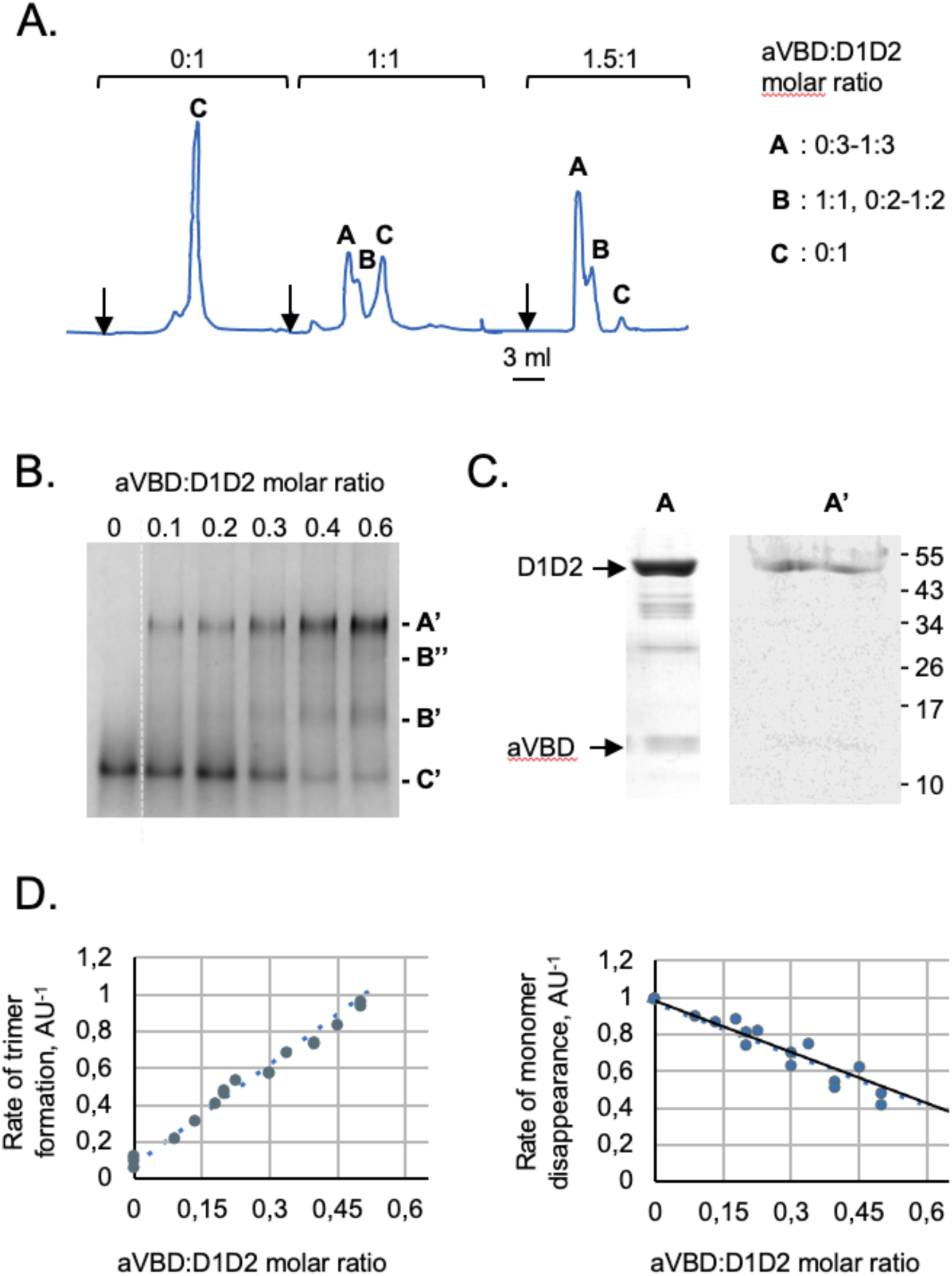
A CN-PAGE assay to study IpaA-induced D1D2 trimer formation. **A**, **B**. aVBD and D1D2 were mixed at the indicated molar ratio, with D1D2 at a final concentration of 20 μM, and incubated for 60 min at 21°C prior to analysis. **A**, SEC analysis using an Increase Superdex 200 (Materials and Methods). The indicated stoichiometry is inferred from previous SEC-MALS analysis (Valencia-Gallardo et al., 2023) molecular and the mass of complexes in peaks A, B and C estimated from molecular weight standards as a function of the respective elution volume. **B**, CN-PAGE using a 7.5 % polyacrylamide native gel followed by Coomassie blue staining. C’: monomeric D1D2.4, B’ and B”: aVBD:D1D2 complexes. **C**, SDS-PAGE followed by Coomassie blue staining of samples corresponding to peak A in SEC fractionation as shown in panel A (A) and eluted following dissection of band A’ as shown in Panel D (A’). Densitometry analysis of the bands indicated a D1D2:aVBD ratio of 3.8 and 3.2 for sample A and A’, respectively, consistent with 3:0-3:1 aVBD complexes. **D**, the integrated intensity of bands corresponding to trimeric (A’) or monomeric D1D2 (C) were scanned by densitometry, and normalized to that of monomeric D1D2 in the absence of aVBD. The graphs are representative of at least three independent experiments.

We used scanning densitometry, to determine the rates of D1D2 trimer formation and D1D2 monomer disappearance normalized to the initial amounts of D1D2 (Materials and Methods). As shown in Fig. 2D, the appearance of D1D2 trimers and disappearance of *apo* D1D2 as a function of increasing aVBD molar ratio could be nicely adjusted to linear fits with a Pearson coefficient R^2^ > 0.95. The CN-PAGE assay was used to determine a rate of D1D2 trimer appearance of 1.51 AU^-1^ ± 0.3 (SEM), and D1D2 monomer disappearance of -1.06 AU^-1^ ± 0.05 (SEM) (Fig. 3A).

**Fig. 3.**
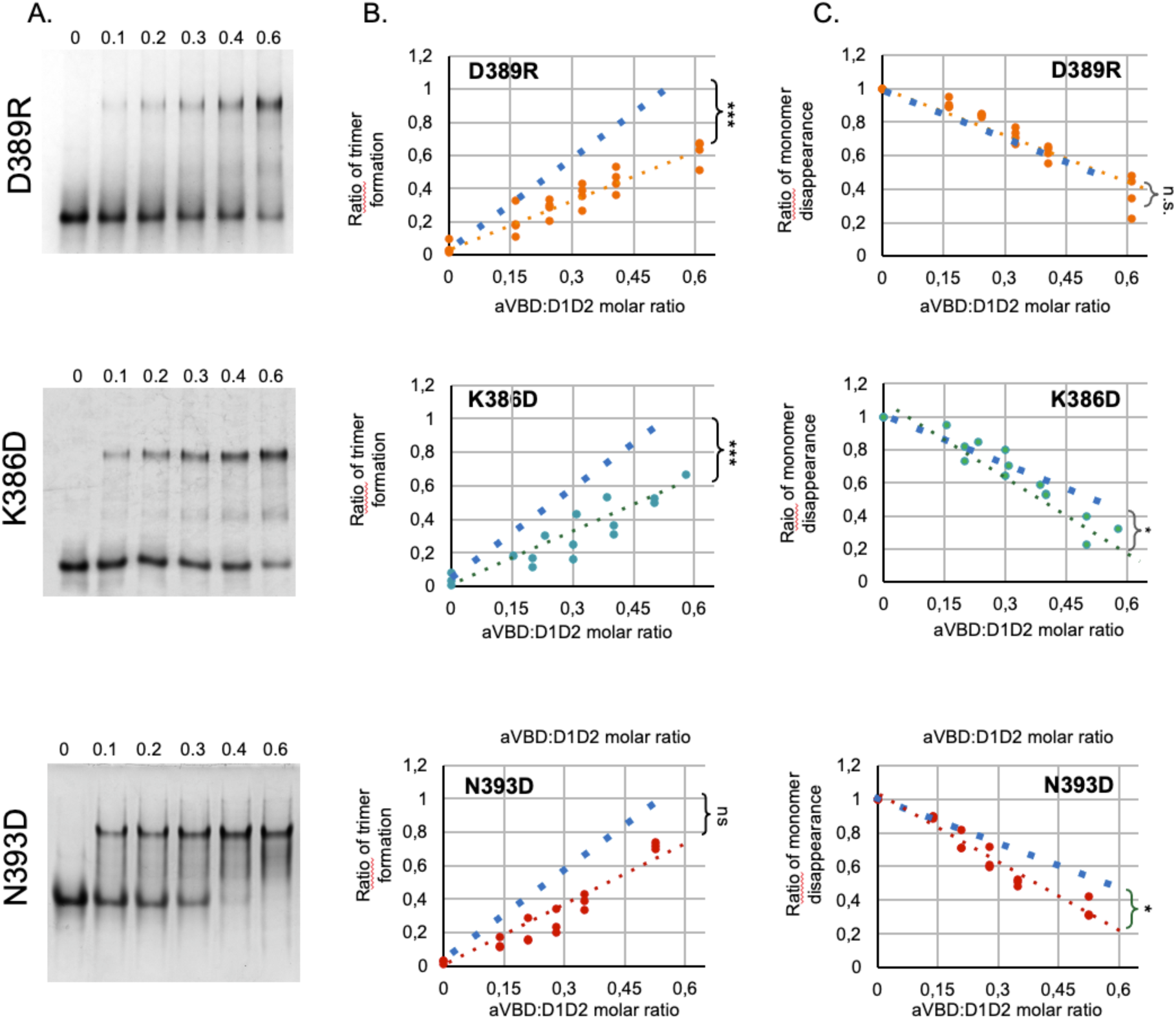
Effects of mutations on D1D2 trimer formation and monomer disappearance. **A**, CN-PAGE using a 10 % polyacrylamide native gel followed by Coomassie blue staining of aVBD-induced complex formation with the indicated D1D2 variant. The aVBD:D1D2 molar ratio is indicated above each lane. **B**, **C**, the integrated intensity of bands corresponding to trimeric (**B**) or monomeric D1D2 (**C**) were scanned by densitometry, and normalized to that of monomeric D1D2 in the absence of aVBD. The graphs are representative of at least three independent experiments. The blue dashed lines correspond to linear fits obtained for parental D1D2. ANCOVA test: *: p < 0.05; ***: p < 0.005; ns: not significant.

### Characterization of D1D2 mutations affecting trimer formation

We estimated that the CN-PAGE assay was sufficiently robust and reproducible to determine potential differences in rates of trimer formation in the various D1D2 variants. All mutations were introduced in D1D2 and the corresponding variants were purified to homogeneity (Fig. S2). Samples were incubated with increasing molar ratio of the aVBD vinculin-binding domain (aVBD) and analyzed by CN-PAGE followed by Coomassie staining (Materials and Methods). As shown in Figs. 3 and S3, the majority of mutants showed decrease rates of trimer formation (Figs. 3A and S3). The rates of D1D2 trimer formation and monomer disappearance were then inferred from linear fits, following quantification by scanning densitometry.

As shown on Figure 3A, all mutations showed reduced rates of D1D2 trimer but could be distinguished in two types based on to their effects on the rates of monomer disappearance. Type 1 mutations D389R, E143K and V367D showed no difference in D1D2 monomer disappearance, while type 2 mutations K386D, K386G and K139E showed increased rates in D1D2 monomer disappearance (Fig. 3A). Mutation N393D showed effects similar to the latter mutations, with a slightly reduced rate of trimer formation that appear not significant, but increased rates in monomer disappearance (Fig. 3A). This 2 types generally correlate with the mutational design based on structural modeling, since type 2 and type 1 mutations are expected to target formation of the closed and open D1D2:aVBD 1:1 complex, respectively (Figs. 1C, D). The D389R mutation represents an exception, since it is expected to target the D1D2:aVBD 1:1 close complex, but showed no difference in the rate of monomer disappearance (Fig. 3A).

In Figure 3B, we depicted potential steps of aVBD-induced D1D2 trimerization affected by the mutations. We posited that type 2 mutations stimulated the transition to the closed 1:1 D1D2:aVBD complex immediately occurring upon aVBD binding to apo D1D2, while inhibiting the subsequent step leading to the open D1D2 conformer (Fig. 3B). In contrast, the type 1 E143E and cysteine clamp mutations that did not affect the rate of monomer disappearance, may target a later step associated with the “opening” of the D1D2 subdomains. The V367D “coiled-coil” mutation could also target a later step impairing D1D2 oligomerization (Fig. 3B). The reasons for the increased rates or absence of effects on monomer disappearance for type 2 mutations and D389R, respectively, while they all target the close D1D2: aVBD conformer, are not clear.

It is possible that type 2 mutations favor the formation of D1D2 dimers that do not trimerize, such as those induced by aVBS1-2 (Valencia-Gallardo et al., 2023). The D389R mutation may mimic the effects of IpaA K499 on the closed D1D2 conformer, therefore explaining the absence of difference on aVBD-induced monomer disappearance, while preventing transition to the open D1D2 conformer (Fig. 3B).

### D1D2 Mutations targeting the close conformer prevent *Shigella* invasion

During *Shigella* invasion, targeting of vinculin by IpaA leads to the cytoskeletal reorganization allowing bacterial attachment to host cells and productive bacterial internalization. In addition, IpaA-mediated vinculin supra-activation triggers the formation of focal adhesions (FAs) distal to the invasion sites that likely contribute to strengthen the adhesion of infected cells even in the absence of mechanotransduction (Valencia-Gallardo et al., 2023). To analyze the role of D1D2 head-domain oligomerization on *Shigella* invasion, D1D2 mutations were transferred to full-length vinculin fused to mCherry (HV-mCherry) cloned in a eukaryotic expression vector and transfected into MEF vinculin -/- cells (Materials and Methods). Transfected cells were challenged with bacteria and bacterial association and internalization were quantified (Materials and Methods).

As shown in Figure 4, the profiles of bacterial association and invasion were generally similar, suggesting consistent with a prominent role in bacterial attachment to the cell during *Shigella* invasion. Consistent for the previously described role for vinculin in *Shigella* invasion, cells complemented with HV-mCherry showed a ca 8-fold and 30-fold increase in bacterial association and invasion compared to mCherry transfected control cells, respectively (Figs. 4B and 4C, HV and mCherry). Similar differences in bacterial association and invasion were observed when comparing to HV-mCherry transfected cells challenged with a non-invasive isogenic mutant *Shigella* strain (Figs. 4B and 4C, HV and BS176). Strikingly, mutations D389R and N393D resulted in a drastic reduction of bacterial association and invasion of host cells, to levels comparable to the negative controls (Figs. 4B and 4C). In contrast, the other tested mutations had modest or non-significant effects. These results suggest a key role for polar and charged interactions involving the D389 and N393 at the N-terminal extremity of the H10 helix of D2. The absence of effects associated with the CC and E143K mutations targeting the open D1D2 conformer suggested that early steps linked to vinculin supra-activation but not later steps linked to vinculin oligomerization are required for *Shigella* invasion.

**Figure 4.**
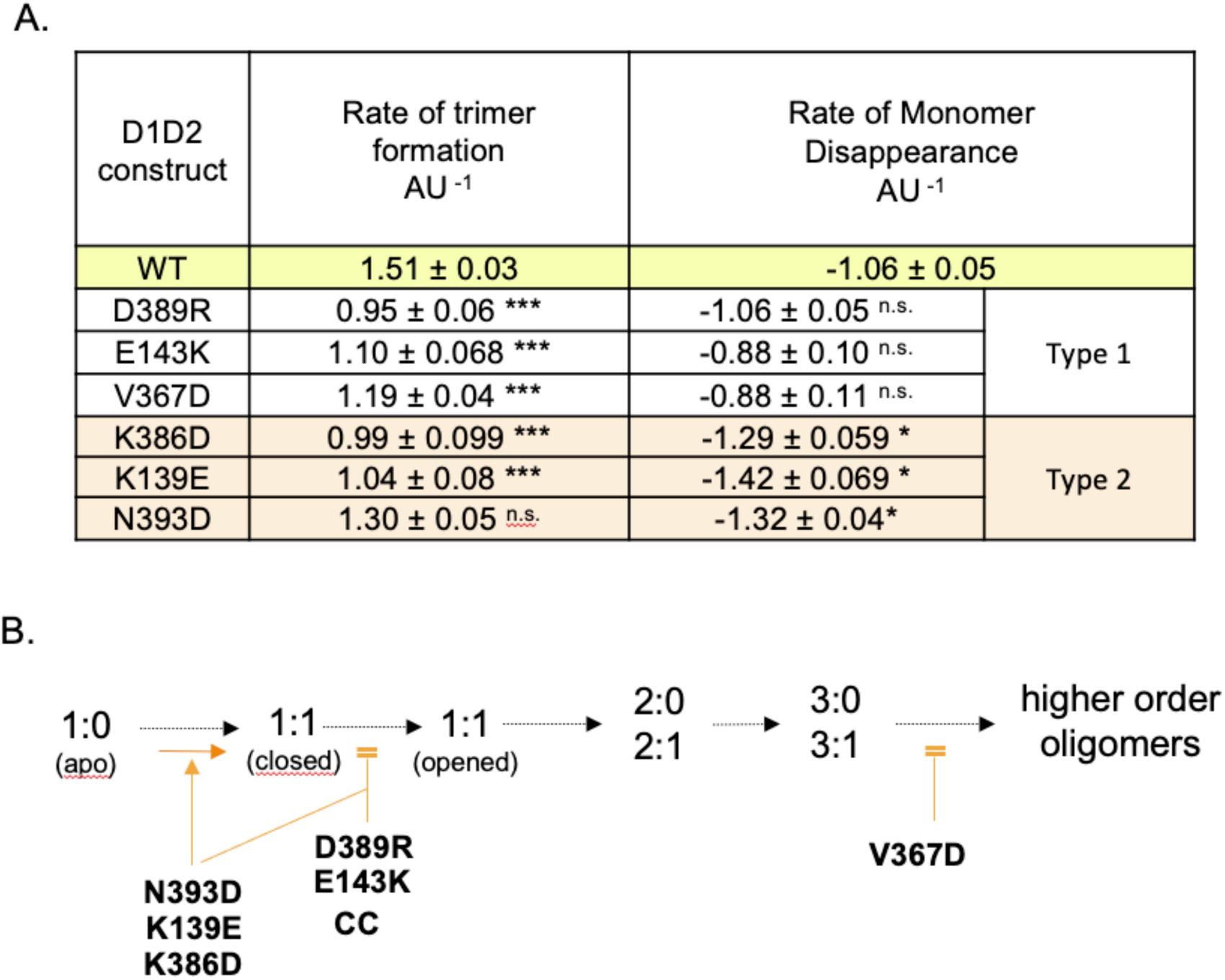
Effects of D1D2 mutations on D1D2 complex formation. **A,** the rates of trimer formation and monomer disappearance for the indicated D1D2 variants were inferred from linear fits (Materials and Methods). Each value corresponds to the mean of at least three independent experiments ± SD. ANCOVA test: *: p < 0.05; **: p < 0.01; ***: p < 0.005. ns: not significant. B, scheme of putative steps in the formation of D1D2:aVBD complexes. The D1D2:aVBD molar ratio is indicated. Orange arrow: mutation favoring complex formation. Orange double bars: mutation impairing complex formation.

### Mutations affecting D1D2 trimerization impair focal adhesions’ formation

Vinculin oligomerization has been mainly studied *in vitro* and reported to occur through the Vt-tail domain (Thomson et al., 2013). In cells, the role of vinculin oligomerization remains unclear. We therefore tested the effects of D1D2 mutations of FA formation, a process relevant for bacterial infection of host cells since it controls the adhesion / detachment of *Shigella*-infected cells from the extracellular matrix.

As shown in Fig. 5, all tested mutations induced a decrease either in FA area or FA number per cell (frequency) or both, although to variable extend (Figs. 5 and S4). The K386D mutation showed a decrease in FA area and frequency but the difference relative to parental HV was not statistically significative, as opposed to the other mutations (Figs. 5B and 5C). As previously reported, the CC mutation altered the size and frequency of FAs (Fig. 5; Valencia-Gallardo et al., 2023). The K139E mutation also significantly affected both FA size and frequency. The E143K, D389R and N393D mutations affected the size of FAs, while the difference in FA number per cell relative to parental HV was not significant because of the large dispersion of values (Figs. 5B and 5C). Interestingly, the V367D variant also did not show a significant difference in the average FA size compared to control cells expressing parental mCherry-vinculin (Figs. 5B and 5C), despite a significant reduction in the number of FAs per cell. The FAs in the V367D variant, although similar in size, looked different than those in control cells, being mostly at the cell periphery and particularly elongated (Fig. 5A). This qualitative difference was confirmed by quantification of the AR index, showing a pronounced difference for the V367D variant compared to the other samples (Fig. S4B).

**Fig. 5.**
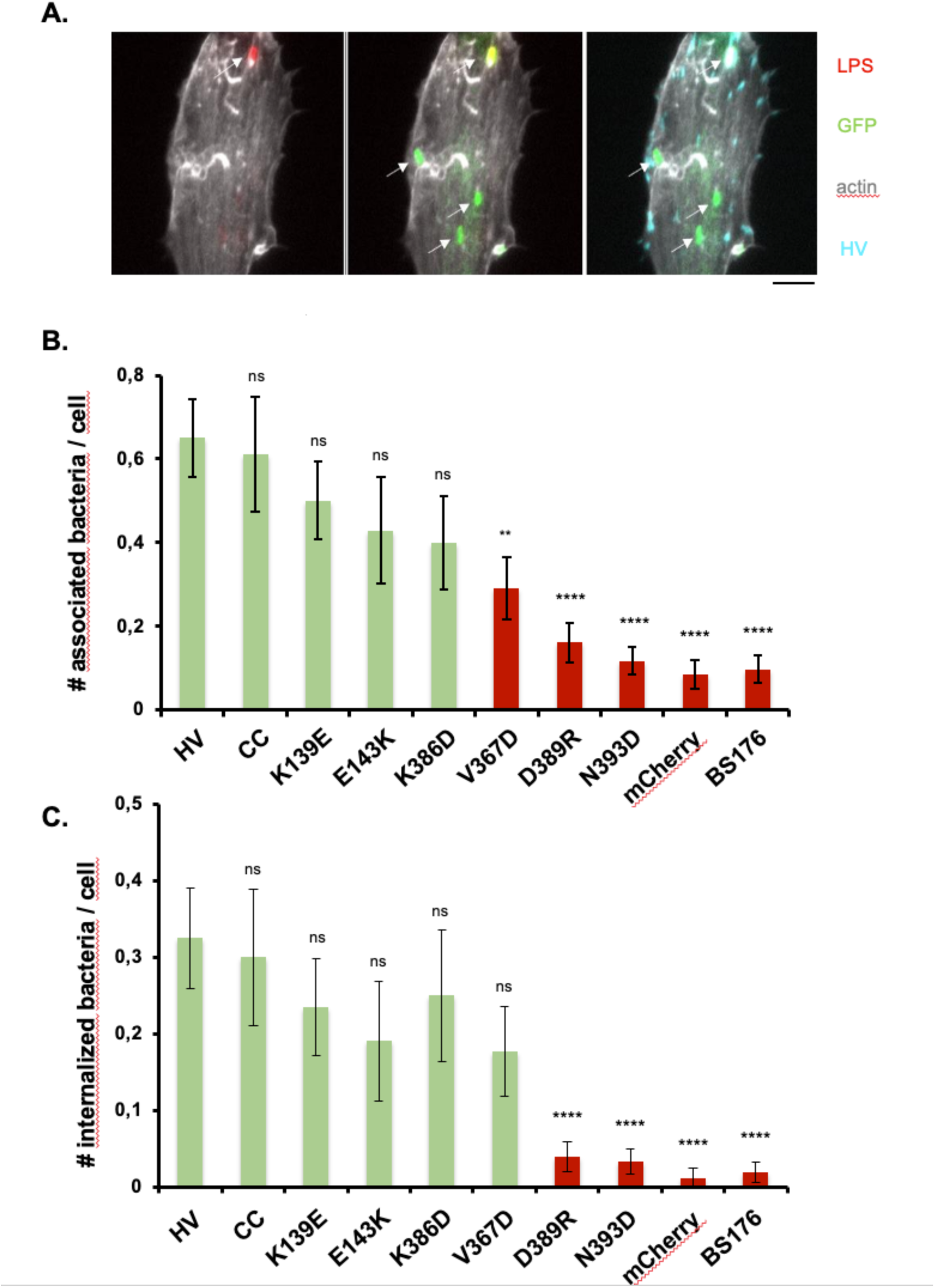
Effects of D1D2 mutations on *Shigella* invasion. MEF vinculin -/- cells were transfected with HV-mCherry (HV) or variants bearing the indicated mutations and challenged with GFP-expressing wild-type *Shigella* for 45 min at 37°C (Materials and Methods). mCherry: cells transfected with mCherry. BS176: cells transfected with HV and infected with or invasion deficient isogenic derivative strain BS176. Following bacterial challenged, samples fixed and processed for bacterial fluorescence inside-out immunostaining and confocal microscopy analysis (Materials and Methods). **A**, representative maximal projection of fluorescent confocal micrographs of cells transfected with HV and challenged with wild-type *Shigella*. Staining: green: GFP; red:LPS; cyan: mCherry; gray levels: F-actin. Scale bar: 5 μm. Note the extracellular bacteria stained in red and green, while internalized bacteria stained in green only. **B**, number of cell-associated bacteria per cell. C, number of internalized per cell. FA area (n > 30 cells, N = 3). **C**, average number of FAs per cell (n > 30 cells, N = 3). T-test: ****: p < 0.001; ns: not significant.

Together, these results support a role for vinculin supra-activation and head-domain mediated oligomerization in the formation of focal adhesions independent of *Shigella* invasion.

### IpaA VBSs prevents cell motility

Our findings suggested that during *Shigella* invasion, IpaA-mediated vinculin head-domain oligomerization played mostly a role in up-regulating cell adhesion rather than bacterial invasion. Vinculin, however, is paradoxically described as a prognostic marker favoring the migration of cancer cells or as a tumor suppressor stimulating cell anchorage (Mierke et al., 2010; Hamidi and Ivaska, 2018). These contradictory findings reflect its complex and poorly understood regulation, as well as different roles in 2D or 3D systems (Gulvady et al., 2018).

We therefore tested the effects of IpaA on the motility and invasion of melanoma cells. In time-lapse microscopy experiments in 2D-chambers, GFP-aVBS1-2 inhibited melanocyte motility compared to control cells, with a rate of Root Median Square Displacement (rMSD) of 3.16 and 15.6 μm.min-1, respectively (Figs. 6A and 6B). An even stronger inhibition was observed for GFP-aVBD transfected cells (rMSD = 2.3 μm.min-1) (Figs. S5A, S5B). Transmigration of melanocytes in 3D-matrigels was similarly inhibited by aVBS1-2 and aVBD (Fig. S5C).

**Fig. 6.**
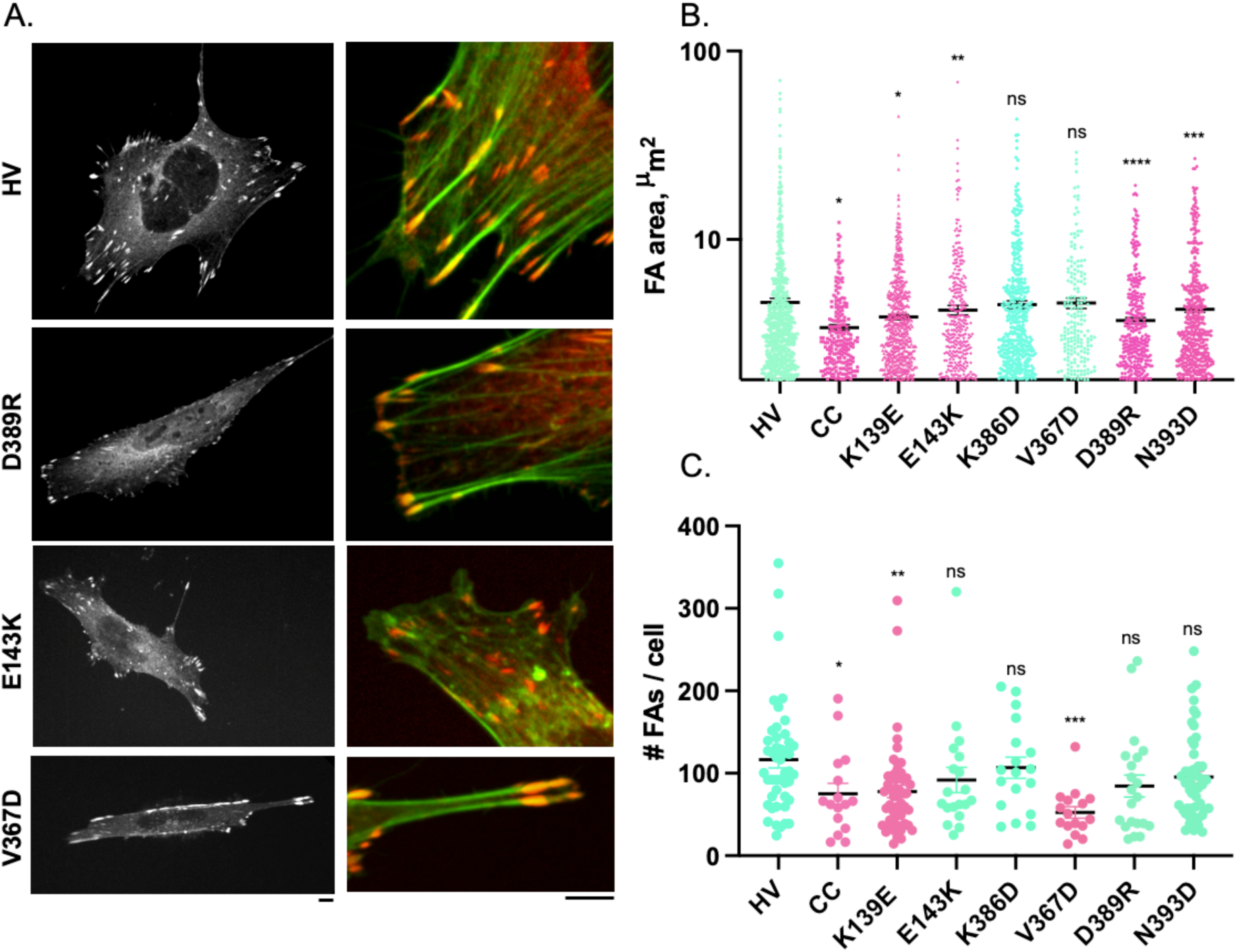
Effects of D1D2 mutations affecting D1D2 trimer formation on focal adhesions. MEF vinculin -/- cells were transfected with HV-mCherry (HV) or variants bearing the indicated mutations. Samples were fixed and processed for fluorescence staining of F-actin. **A**, representative maximal projections of fluorescent confocal micrographs. Green: F-actin; red:HV-mCherry. Scale bar: 5 μm. **B**, FA area (n > 1500, N = 2). **C**, average number of FAs per cell (n > 20, N = 2). Mann-Whitney: *: p < 0.05; **: p < 0.01; ***: p < 0.005. ns: not significant.

**Figure 7.**
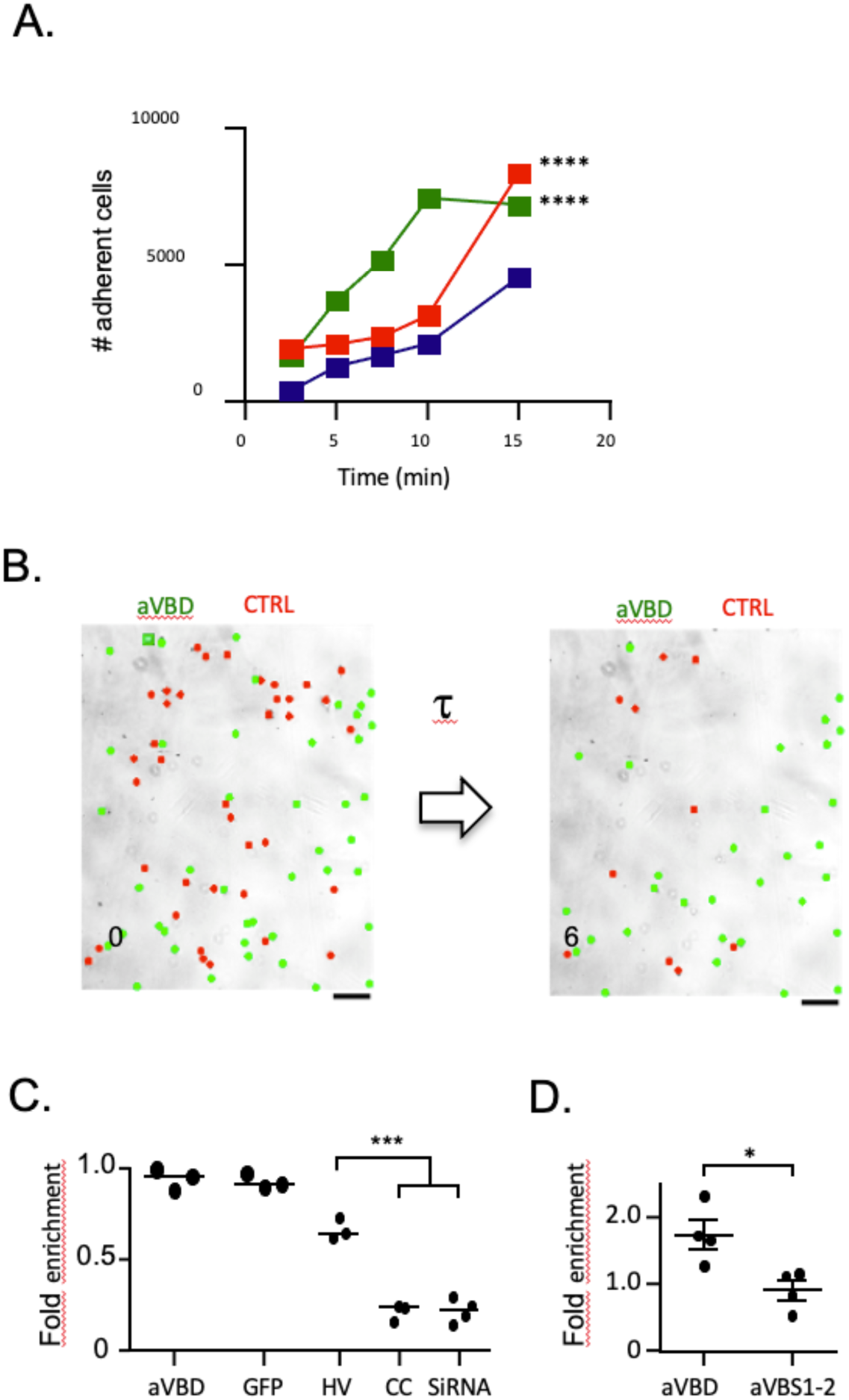
aVBD-mediated vinculin supra-activation stimulates rapid cell adhesion. **A**, 1205Lu melanoma cells were transfected with GFP alone (blue), GFP-aVBS1-2 (red) or GFP-aVBD (green), lifted up by trypsinization and plated for the indicated time on Fn-coated coverslips. Samples were washed, fixed and adherent cells were scored microscopically. The total number of adherent cells scored is indicated. GFP: 3223 cells, N = 4; GFP-aVBS1-2: n=7418, N = 4; GFP-aVBD: n = 5668, N = 4. Chi square corrected with Bonferroni mutliple comparison correction. ****: p < 0.05. **B-D**, 1205Lu melanoma cells were transfected with the indicated constructs, labeled with calcein (methods) and mixed with the same ratio of control cells. Cells were perfused in a microfluidic chamber and allowed to adhere for: **C**, 20 min; or **D**, 15 and 30 min prior to shear stress application. **B**, representative fields. The number indicates the elapsed time after shear stress application. **C**, **D**, Scatterplot of fold enrichment of: aVBD (N = 4, n = 610) or aVBS1-2 (N = 4, n = 433) transfected cells vs control cells (1594 cells, N = 4). **D**, Cells were allowed to adhere for 30 min (large solid circles) or 15 min (small solid circles) prior to shear stress application. aVBD (N = 3, n = 557); GFP (N = 3, n = 490); HV: vinculin mCherry (N = 3, n = 481); CC: vinculin Q68C A396C-mCherry (N = 3, n = 259); siRNA: cells treated with anti-vinculin siRNA (N = 3, n = 395). Unpaired t test. ***: p < 0.005.

These results suggest that IpaA-mediated vinculin “canonical” activation induced by aVBS1-2, as well as “supra-activation” mediate by aVBD do not stimulate cell motility but rather strengthen cell adhesion, consistent with their described effects of FAs (Valencia-Gallardo et al., 2023).

### Vinculin supra-activation, catalyzed by IpaA, strengthens cell adhesion

We next investigated the role of vinculin supra-activation by comparing the effects of aVBS1-2 and aVBD in dynamic cell adhesion experiments. First, we performed cell replating experiments, where resuspended transfected cells were allowed to attach to fibronectin-coated surfaces for controlled time periods. As shown in Fig. 6A, aVBD induced higher yields of adherent cells when replating was performed with short kinetics, with a 5-fold increase over control cells or cells transfected with aVBS1-2 after 10 min replating. By contrast, little difference in adhesion yield was detected between samples after 15 min replating suggesting that aVBD predominantly affected the early dynamics rather than increasing cell adhesion (Fig. 6A). To extend these findings, we measured cell adhesion strength using controlled shear stress in a microfluidic chamber (Fig. 6B; Suppl. movie 1). Consistent with replating experiments, when cells were allowed to adhere to fibronectin-coated surfaces for more than 25 min, little difference in resistance to shear stress could be detected between GFP-aVBD and GFP transfected cells samples (Fig. 6C). In contrast, similar to cells depleted for vinculin by siRNA treatment, cells transfected with the clamped vinculin version showed a decreased ability to adhere in comparison to wild-type vinculin-transfected cells (Figs. 6C and S5). However, when shear stress was applied after less than 20 min following cell incubation, GFP-aVBD-transfected cells showed significantly higher resistance to shear stress up to 22.2 dynes.cm-2 compared to GFP-aVBS1-2 or GFP-transfected cells, with 1.7 ± 0.2 -and 0.9 ± 0.14-fold enrichment ± SD of adherent cells for GFP-aVBD and GFP-aVBS1-2 or GFP transfected cells, respectively (Fig. 6D; Suppl. movie 1). These results are in full agreement with effects observed on adhesion structures and suggest that aVBD-mediated vinculin supra-activation accelerates endogenous processes occurring during mechanotransduction to promote strong adhesion.

## Discussion

In this work, we identified polar residues located at the C-terminal extremity of the H10 helix in vinculin D2 involved in D1D2 trimerization induced by *Shigella* IpaA. Among mutations reducing IpaA-induced D1D2 trimer formation *in vitro*, we identified the charge inversions at vinculin D389R and E143K that do not affect the rates of monomer disappearance, while the K386D polar mutations led to increased rates of monomer disappearance. IpaA vinculin interaction is mediated by 3 VBSs, IpaA VBS1 and VBS2 interacting with the first and second bundle of D1, respectively. The high affinity of aVBS1-2 to D1 is likely to drive the initial steps leading to the formation of a D1D2: aVBD 1:1 complex, independent of IpaA VBS3. From our structural models of the D1D2:aVBD 1:1 complexes, the vinculin D389R and E143K mutations are expected to disfavor IpaA VBS3 binding to the close and open D1D2 conformer, respectively. Hence, the effects of these mutations are in line with specific effects on IpaA VBS3 allosteric changes leading to D1D2 oligomerization without affecting formation of aVBS1, 2-dependent 1:1 D1D2:aVBD complex.

There are two possible explanations to the effects of the K139E, K386D and N393D mutations that decrease the rates of D1D2 trimer formation while increasing monomer disappearance: i, these polar mutations may increase the rates of IpaA binding to D1D2. This possibility is unlikely because of the predominant role of aVBS1-2 in driving the formation of a D1D2:aVBD 1:1 complex and because these mutations were precisely designed to interfere with IpaA VBS3 interaction with D1D2; ii, they may stabilize an allosteric D1D2 conformer subsequent to the formation of the D1D2:aVBD 1:1 complex, thereby favoring the formation of this latter complex and accelerating monomer disappearance. This possibility is counter-intuitive because it implies that these polar mutations have opposite effects on favoring the formation of an D1D2:aVBD intermediate complex while disfavoring the formation of the trimer. However, it needs to be envisioned since the IpaA VBS3 helix is unlikely to function in the same iterative manner during D1D2 dimerization and resolution into a D1D2 homotrimer, and rather interacts with different subset of residues during these processes. The effects of the K139E mutation also impairing D1D2 trimer formation and accelerating monomer disappearance may be explained in a related yet different manner. Indeed, in apoD1D2, K139 establish a salt bridge with D389 that likely stabilize the D1D2 in the close conformation. The vinculin K139E charge inversion may therefore destabilize D1D2 to favor the formation of intermediate D1D2:aVBD complexes.

We found that the V367D predicted to disrupt the putative coiled-coil heptad motif in D2 H10 affected trimer formation, suggesting a role for this motif in IpaA-induced D1D2 oligomerization. In the structures of apo D1D2 of full-length vinculin, this motif is located at a face of H10 buried into the D2 second helix bundle both in the apo/closed and open D1D2 conformer. Exposure of this putative coiled-coil motif would require major unraveling of the D2 bundles and high pulling force ranges. Using atomic force microscopy and steered molecular dynamics, Kluger et al. estimated that pulling forces ranging from 30-60 pN could lead to major rearrangements of the D1 with some unraveling of the second D1 bundle, while no such reorganization was observed for the other vinculin head subdomains (Kluger et al., 2020). These measurements, however, were performed at a single vinculin molecule level. I Different D1D2 interfaces under a vinculin trimeric form combined with higher exerted forces expected from actin filaments’ bundling may result in D2 unfolding, leading to exposure of the coiled-coil motif and vinculin higher order oligomerization (Fig. 4B).

Our findings based on a clamped-mutant of vinculin proficient for canonical activation but deficient for supra-activation suggest that the head-domain mediated oligomerization of vinculin akin to that induced by IpaA is required for the maturation of adhesions into large focal adhesions (Valencia-Gallardo et al., 2023). The observed effects of the D1D2 mutations support this hypothesis, since these mutations affecting vinculin trimerization introduced into full-length vinculin affected the number and size of on focal adhesions. While showing reduced and smaller focal adhesions, the amplitude of these defects was less in the K386D vinculin variant. These findings may be related to the different effects of this mutation observed *in vitro* relative to the D389R and E143K mutations, and to the accumulation of intermediate vinculin oligomeric complexes. Indeed, we previously found that in replating experiments, IpaA favored the rapid adhesion of cell to the substrate, but that similar levels of adhesion were detected over prolonged incubation upon canonical activation including that induced by an IpaA variant deleted for VBS3 (Valencia-Gallardo et al., 2022). The findings suggested that IpaA could promote the supra-activation of vinculin in the absence of mechanotransduction, but that canonical activation associated with mechanotransduction could also induce supra-activation. It is possible that the accumulation of oligomeric complexes linked to the K386D mutation compensate the defect of IpaA-induced trimers, during mechanotransduction.

The size reduction of focal adhesions associated with D1D2 mutations was more specifically observed at the levels of peripheral adhesions, while little difference was observed for ventral adhesions relative to parental vinculin. Ventral adhesions appeared thin and elongated, consistent with fibrillar adhesions known to remain stable independent of force (Sun et al., 2016). These observations are consistent with head domain-mediated vinculin oligomerization occurring at high force regime. As opposed to other mutations, the V367D mutation led to a drastic reduction of the number of focal adhesions but did not affect their average size. Instead, the V367D variant mostly formed elongated adhesions at the cell periphery. While the role of the putative coiled-coil domain targeted by the V367D mutation deserves clarification, the effects on the focal adhesion morphology associated with this particular mutation is consistent with the impairment in a process during vinculin oligomerization and focal adhesion formation that is different than the other mutations.

We found that as opposed to the effects observed for all D1D2 mutations on focal adhesions, only the N393D and D389R mutations affected *Shigella* invasion of host cells. These mutations are predicted to impair the interaction of IpaAVBS3 in the closed but not in the open D1D2:aVBD conformer. Since in this closed conformer, D1D2 adopt a conformation similar to that of D1D2:aVBS1-2 or apo D1D2, this suggests that it occurs prior to formation of the open D1D2:aVBD conformer that leads to vinculin head-domain oligomerization (Fig. 4B; Valencia-Gallardo et al., 2023). In contrast, D1D2 mutations targeting the open conformer and vinculin oligomerization had no significant effects on bacterial invasion. The results suggests that IpaA-mediated vinculin supra-activation and head-domain oligomerization is not critical for *Shigella* invasion, but is required to strengthen matrix adhesion of bacteria-infected cells. Accordingly, rather than acting at the bacteria-cell contact sites where it is injected by the *Shigella* T3SS, IpaA would induce the formation of vinculin oligomers diffusing at a distance from the supra-activation site to reinforce cell adhesions at basal membranes. This view is in line with the “catalytic” action of IpaA on vinculin oligomers formation observed *in vitro* (Valencia-Gallardo et al., 2023).

## Materials and Methods

### Bacterial strains, cells and plasmids

The bacterial strain used for the purification of D1D2 construct is *E. coli* BL21 (DE3) from Invitrogen. *E. coli* DH5-α F– *endA1 glnV44 thi-1 recA1 relA1 gyrA96 deoR nupG purB20 φ80dlacZΔM15 Δ(lacZYA-argF) U169, hsdR17(rK–mK+), λ– was* used for the purification of aVBD. MEF and MEF vinculin null cells (Humphries, Wang et al. 2007) were grown in DMEM 1 g / L glucose containing 10 % FCS in a 37°C incubator containing 10 % CO_2_.

The pGEX4T2-aVBD and the pET15b-D1D2 plasmids were described previously. The mutations in D1D2 were introduced by site-directed mutagenesis using pET15b-D1D2 as a matrix and the primer pairs indicated in Table 1. The pmCherry-N1-human vinculin (HV) and was from Addgene. Mutations in D1D2 were transferred in mCherry-HV by exchanging the NheI-PspXI fragment with the corresponding XbaI-PspXI fragment of pET15b-D1D2. MEF vinculin null cells (Humphries, Wang et al. 2007) were routinely grown in DMEM 1 g / L glucose containing 10 % fetal calf serum in a 37°C incubator containing 10 % CO_2_.The 1205Lu melanoma cell line, a gift from Dr. M. Herlyn (Wistar Institute, Philadelphia, PA), was cultured in RPMI supplemented with 10% fetal calf serum as described (Javelaud et al. 2005).

### Protein purification

BL21 (DE3) competent *E. coli* was transformed with the pET15b-D1D2 variant constructs. D1D2 were purified as described (Park, Valencia-Gallardo et al. 2011). For the IpaA derivatives, DH5-a competent *E. coli* was transformed with pGEX-4T2-aVBD. Bacteria were grown at 37°C with shaking until OD_600nm_ = 1.0 were induced with 1 mM IPTG and incubated for another 2 hrs. Bacteria were pelleted and washed in ice-cold lysis buffer containing 25 mM Tris PH 7.4, 100 mM NaCl and 1 mM beta-mercaptoethanol, containing Complete^TM^ protease inhibitor. All subsequent steps were performed at 4°C. Bacterial pellets were resuspended in 1/20th of the original culture volume and lyzed using a high pressure homogeneizer (LM20, Microfluidics Corp., MA, USA). Cell debris were pelleted by centrifugation at 8000 xg for 20 min. Clarified lysates were subjected to affinity chromatography using a GSTrap HP affinity column (GE Healthcare). Briefly, following incubation with the clarified lysates, the column was washed with five column volumes prior to incubation in PBS containing 100 μg / ml Thrombin (Cytiva, ref 27084601) for 16 hours at 21°C. aVBD was then eluted in PBS and further subjected to purification using size exclusion chromatography using a Superdex 200 10/300 GL (Ge Healthcare). Samples were stored aliquoted at -80°C at concentrations ranging from 1 to 10 mg/ml.

### SEC analysis

D1D2 and aVBD were at the indicated molar ratio, with D1D2 at a final concentration of 20 μM, for 60 minutes at 21°C. 200μl of the protein mixtures were analyzed by size-exclusion chromatography (SEC) on a Superdex 200 10/300 GL (GE Healthcare) using a GE ÄKTA FPLC™ (Fast Protein Liquid Chromatograph, GMI) and a collection volume of 200 μl per fraction and 20ml of total collected volume. The SEC buffer was 25 mM Tris-HCl pH 7.2, 100 mM NaCl.

### CN-PAGE and densitometry analysis

The D1D2 variants and aVBD were at the indicated molar ratio, with D1D2 at a final concentration of 20 μM, for 60 minutes at 21°C. Protein complex formation was visualized by PAGE under non-denaturing conditions using à 7.5% polycrylamide gel, followed by Coomassie blue staining, as described previously (Wittig and Schägger, 2005).

Gel raw images were aquired with a ChemiDoc™ Imaging System (Bio-Rad Laboratories, Inc. USA) using the Coomassie staining settings. Band intensities corresponding to the monomer, higher order oligomer or intermediate species were measured with Fiji-ImageJ using a fixed rectangular area adjusted with the max width for the monomer and max high order oligomer for the height (Schindelin et al., 2019). A reference area outside the lanes used for the running reactions was included to subtract the background. Values were plotted and then normalized to the value obtained for the corresponding apo construct.

### Cell transfection

For transfection experiments, cells were seeded at 1 x 10^4^ cells on 25 mm-diameter coverslips coated with fibronectin at a concentration of 20 μg / ml. Cells were transfected with 1 μg of the pGEX-4T2-D1D2 construct and 4 μls JetPEI transfection reagent (Polyplus) for 16 hours following the manufacturer’s recommendations.

### Fluorescence confocal microscopy analysis

Samples were fixed in PBS containing 3.7 % paraformaldehyde for 60 min at 21°C, prior to processing for fluorescence staining of F-actin using Phalloidin-Alexa 488 as previously described (Valencia-Gallardo et al., 2022). Samples were analyzed using an Eclipse Ti inverted microscope (Nikon) equipped with a 60 x objective APO TIRF oil immersion (NA: 1.49), a CSU-X1 spinning disk confocal head (Yokogawa), and a Prime 95B sCMOS camera (Photometrics) controlled by the Metamorph 7.7 software.

### Image analysis

Focal adhesions were analyzed using the ImageJ 2.1.0/1.53c software. For each set of experiments, the confocal plane corresponding to the basal plane was subjected to thresholding using strictly identical parameters between samples. Adhesion clusters were detected using the “Analyze particle” plug-in, setting a minimal size of 3.5 μm^2^.

### Statistical analysis

The number of adhesions was analyzed using Dunn’s multiple comparisons test. The median area was compared using Mann-Whitney test. Differences in the rates of D1D2 trimer formation and monomer disappearance based were analyzed using an ANCOVA test.

### Live cell tracking

1205Lu melanoma cells were transfected with IpaA constructs or GFP alone (control) and transferred in microscopy chamber on a 37°C 5%-CO_2_ stage in RPMI1640 medium containing 25 mM HEPES. For cell tracking, samples were analyzed using and inverted Leica DRMIBe microscope and a 20 X phase contrast objective. Image acquisitions were performed every 3 min for 200 hrs. The mean velocity of migration was measured for all tracks followed for at least 5 hours. The root square of MdSD over time was plotted over time and fitted by linear regression. The slopes of the linear fit were compared using an ANCOVA test (linear model). The median cell surface was quantified as the mean of the surface for three time points (25%, 50% and 75%) of the whole cell track and dispersion measured by the Median absolute dispersion (MAD).

### Invasion assays

Tissue culture Transwell inserts (8 μm pore size; Falcon, Franklin Lakes, NJ) were coated for 3 hours with 10 μg of Matrigel following the manufacturer’s instructions (Biocoat, BD Biosciences, San Jose, CA). Inserts were placed into 24-well dishes containing 500 μl of RPMI medium supplemented with 1% fetal calf serum. 5 × 10^4^ melanoma cells were added to the upper chamber in 250 μls of serum-free RPMI medium. After 24 hours, transmigrated cells were scored by bright field microscopy. Experiments were performed at least three times, each with duplicate samples.

### Microfluidics cell adhesion assay

Analysis of cell detachment under shear stress was based on previous works [38]. 1205Lu melanocytes were transfected with the indicated constructs, then labeled with 2 μIll calcein-AM (Life Technologies) in serum-free DMEM for 20 minutes. Cells were detached by incubation with 2 μIll Cytochalasin D (Sigma-Aldrich) for 40 minutes to disassemble FAs, followed by incubation in PBS containing 10 mM EDTA for 20 minutes. Cells were washed in EM buffer (120 mM NaCl, 7 mM KCl, 1.8 mM CaCl_2_, 0.8 mM MgCl_2_, 5 mM glucose and 25 mM HEPES at pH 7.3) by centrifugation and resuspended in the same buffer at a density of 1.5 x 10^6^ cells/ml. Calcein-labeled transfected cells and control unlabeled cells were mixed at a 1:1 ratio and perfused onto a 25 mm-diameter glass coverslips (Marienfeld) previously coated with 20 μg/ml fibronectin and blocked with PBS containing 2% BSA (Sigma-Aldrich) in a microfluidic chamber on a microscope stage at 37°C. We used a commercial microfluidic setup (Flow chamber system 1C, Provitro) and a Miniplus3 peristaltic pump (Gilson) to adjust the flow rate in the chamber. Microscopy analysis was performed using a LEICA DMRIBe inverted microscope equipped with a Cascade 512B camera and LED source lights (Roper Instruments), driven by the Metamorph 7.7 software (Universal imaging). Cells were allowed to settle for the indicated time prior to application of a 4 ml/min, flow corresponding to a wall shear stress of 22.2 dyn/cm^2^ (2.22 Pa). Acquisition was performed using a 20 X objective using phase contrast and fluorescence illumination (excitation 480 ± 20 nm, emission 527 ± 30 nm). Fluorescent images were acquired before and after flushing to differentiate between target and control cells. Phase contrast images were acquired every 200 ms. Fold enrichment was defined as the ratio between of attached labeled and unlabeled cells.

## Acknowledgements

The research was supported by fundings from the Inserm and CNRS. GTVN is a recipient of the grant ANR-21-CE35-0007-03 CALPLYCX. DA and BCC received funding from the CONACYT. HK was supported by the French Agence Nationale de la Recherche (ANR), under grant ANR-22-CPJ2-0075-01.

## Author contribution

BCC, HK, MK, DIA, CB, CVG, YZ, DJ and GTVN performed experiments. HK performed the structural modeling. BCC, HK, MK, DIA AND CVG analyzed data. BCC and GTVN conceptualized experiments and wrote the original draft of the manuscript. JC and AM supervised the section related to the cell adhesion in microfluidics experiments and melanocytes invasiveness, respectively. GTVN acquired fundings.

## Supplementary Materials

**Figure S1.**
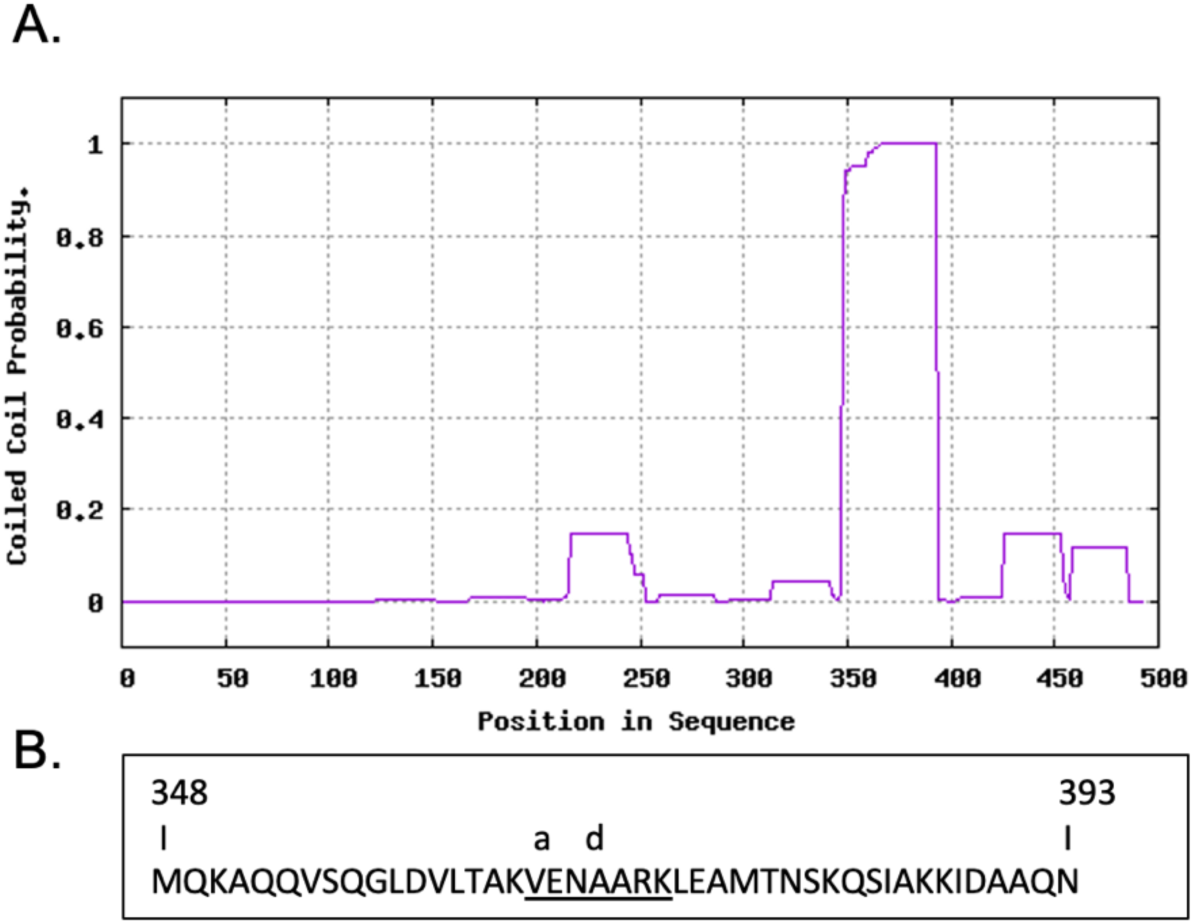
Predicted coiled-coil motif in D2. **A**, the coiled-coil probability in D1D2 corresponding to vinculin residues 1-484 assessed using the MARCOIL algorithm with threshold 50: 11 (Delorenzi and Speed, 2002), identified a coiled-coil region between residues 348-393 with max = 99.9. **B**, sequence of the predicted coiled-coil region identified in A. The numbers correspond to the residue numbers in the vinculin sequence. The canonical heptad region is underlined, with a and d corresponding to hydrophobic residues at position V367 and A370, respectively. The V367D mutation is predicted to abolish coiled-coil formation.

**Figure S2.**
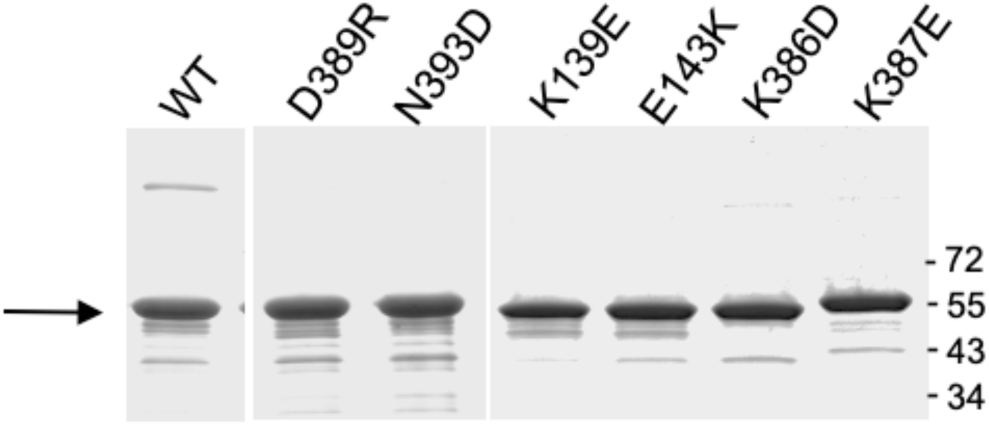
SDS-PAGE analysis of D1D2 variants. D1D2 variants expressed in *E. coli* BL21 (DE3) and purified by affinity chromatography (Materials and Methods) were analyzed by SDS-PAGE using a 10 % polyacrylamide gel followed by Coomassie staining. The size of the molecular weight markers is indicated in kDa. The arrow points at D1D2 constructs.

**Figure S3.**
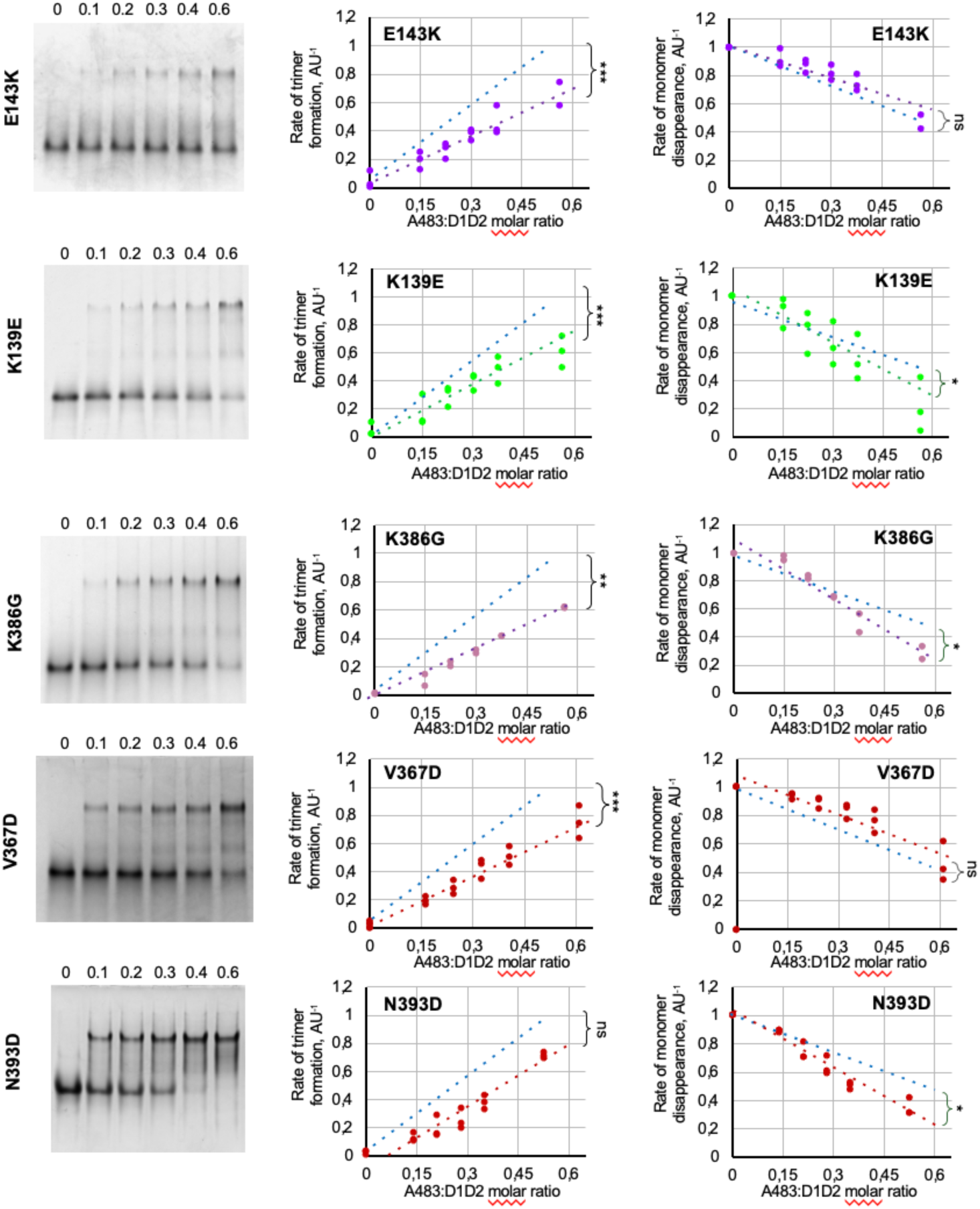
Effects of D1D2 mutations on IpaA-induced trimer formation and monomer disappearance. **Left panels**, CN-PAGE using a 10 % polyacrylamide native gel followed by Coomassie blue staining of aVBD-induced complex formation with the indicated D1D2 variant. The aVBD:D1D2 molar ratio is indicated above each lane**. Center and right panels**, the integrated intensity of bands corresponding to trimeric (**center**) or monomeric D1D2 (**right**) were scanned by densitometry, and normalized to that of monomeric D1D2 in the absence of aVBD. The graphs are representative of at least three independent experiments. The blue dashed lines correspond to linear fits obtained for parental D1D2. ANCOVA test: *: p < 0.05; **: p < 0.01; ***: p < 0.005. ns: not significant.

**Fig. S4.**
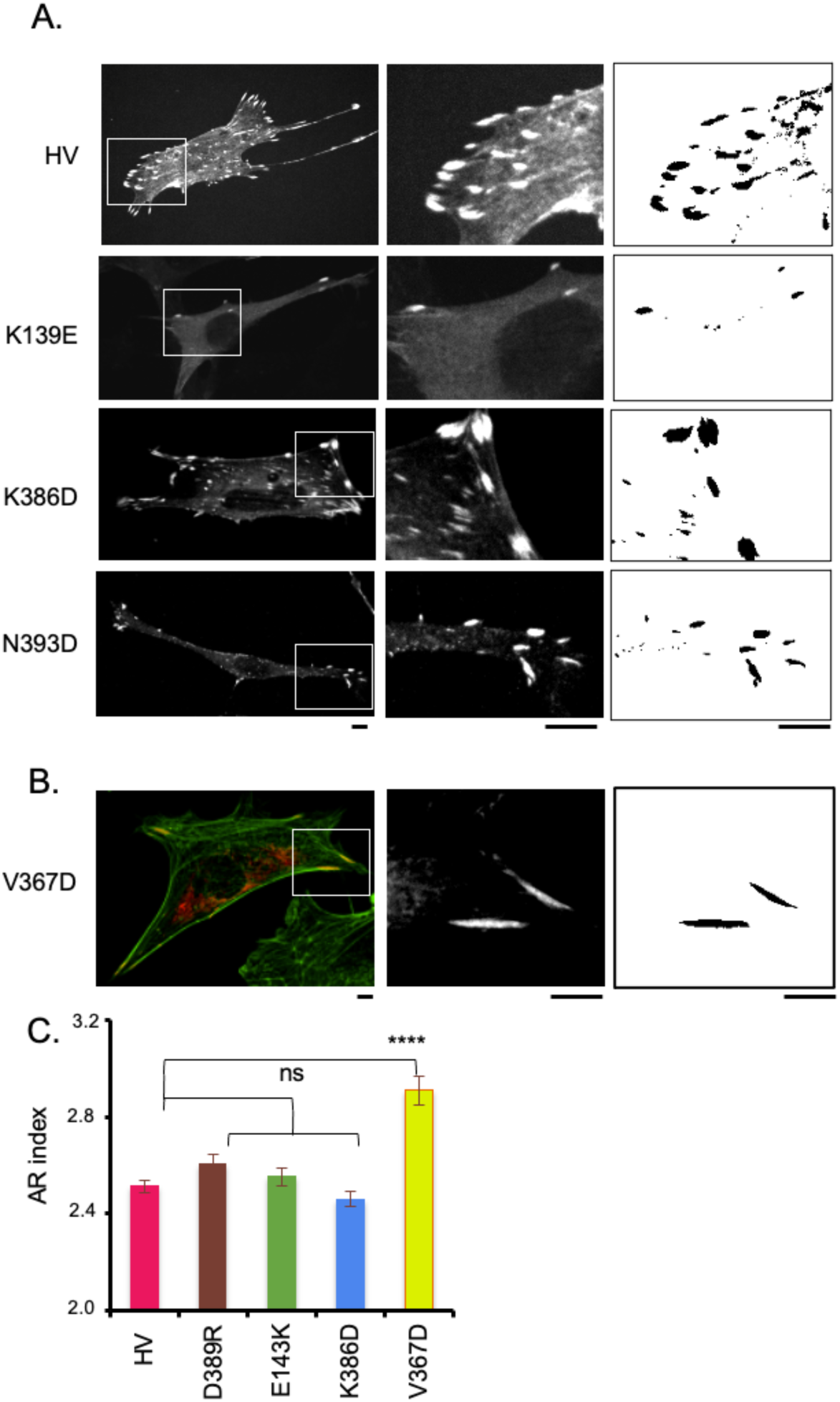
Effects of D1D2 mutations affecting D1D2 trimer formation on focal adhesions. MEF vinculin -/- cells were transfected with HV-mCherry (HV) or variants bearing the indicated mutations. Samples were fixed and processed for confocal fluorescence microscopy analysis of: **A**, mCherry (A); **B**, mCherry (red) and F-actin (green). **A**, **B**, Left: representative maximal projections of mCherry fluorescence. Middle: higher magnification of the corresponding inset shown in left panel. Right: FA detection from overlay masks (Materials and Methods). Scale bar: 5 μm. **C**, AR index corresponding to the major over minor axis of FA. (n > 2500, N = 2). Mann-Whitney: ****: p < 0.001. ns: not significant.

**Figure S5.**
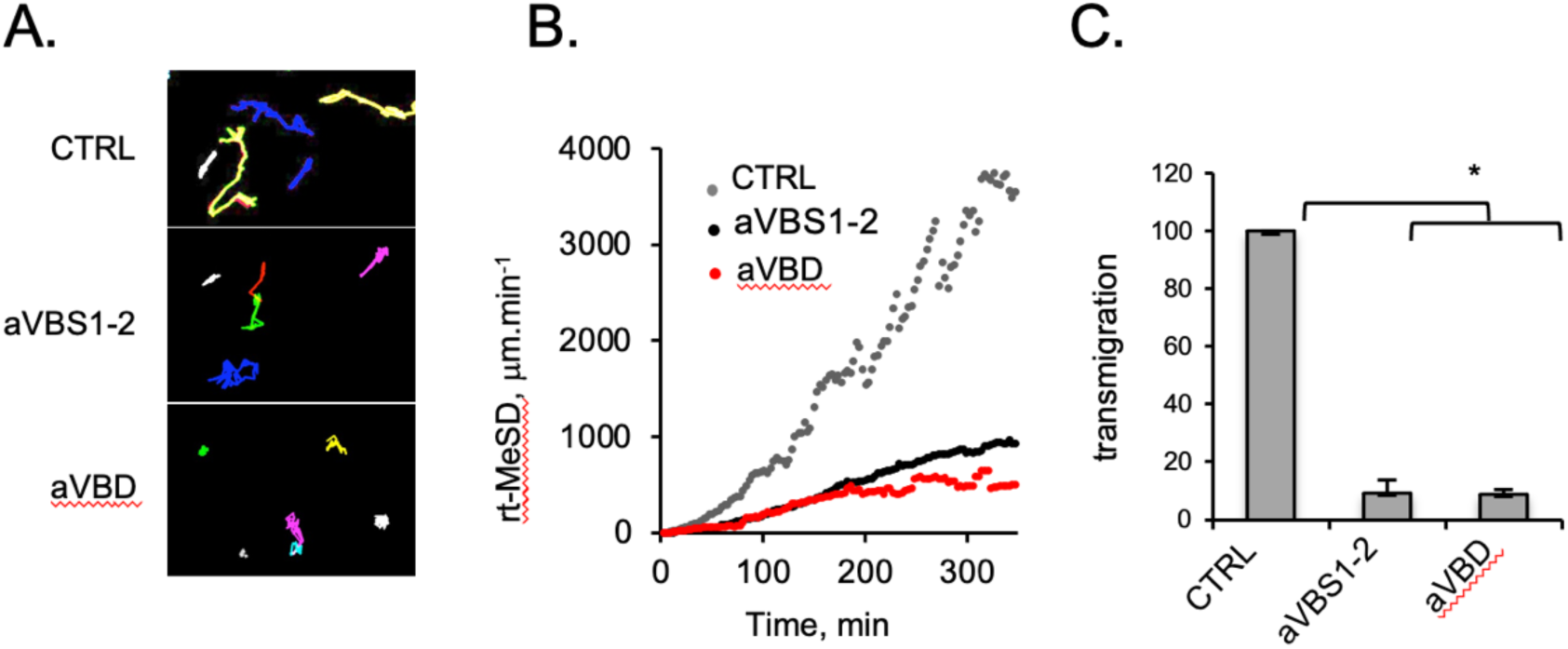
aVBD inhibits tumor cell invasion. 1205Lu melanoma cells were transfected with GFP alone (CTRL), GFP-aVBS1-2 or GFP-aVBD as indicated lifted up by trypsinization and plated for the indicated time on Fn-coated coverslips. Samples were transfected with the indicated constructs, and analyzed by time-lapse videomicroscopy; **A**, representative single cell migration 20-hour tracks for indicated samples. **B**, Root of Median Square of displacement over time for control- (61 cells, N = 3), GFP-A524- (61 cells, N = 3) and GFP-A483 transfectants (64 cells, N = 3). ***: p = 0.0007. The slopes were analyzed using a covariance test and found to be statistically different (ANCOVA, p < 2×10^-16^). **C,** 5 × 10^4^ cells were seeded in matrigel chambers. The percent of transmigrated cells is indicated. (N = 3). Kruskal-Wallis test with Dunn’s multiple comparisons test. *: p < 0.05.

**Figure S6.**
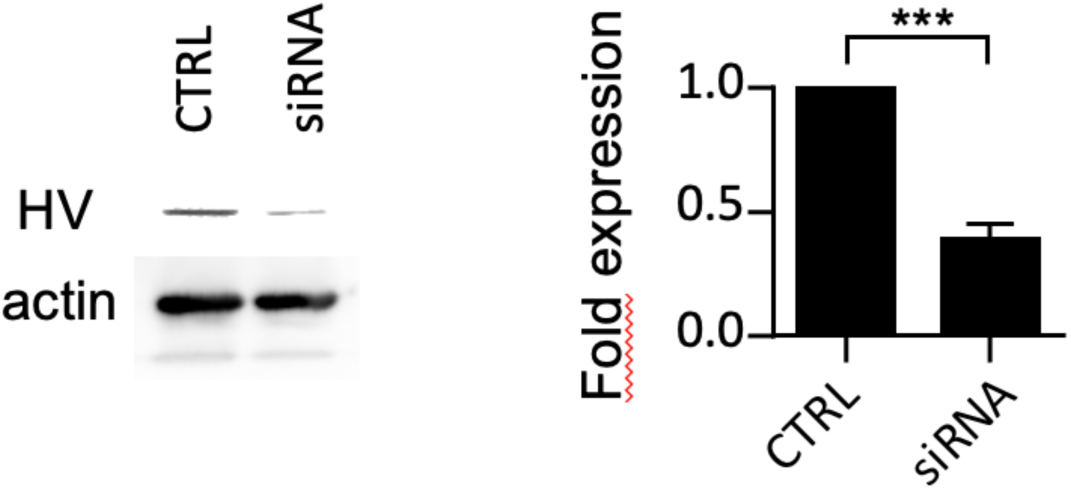
siRNA-mediated inhibition of vinculin expression in 1205Lu melanoma cells. 1205Lu melanoma cells were mock-transfected (CTRL) or treated with anti-vinculin siRNA (siRNA, Methods). **b**, anti-vinculin Western blot analysis. **c**, average HV band intensity normalized to that of control cells. Unpaired t test. ***: p = 0.005.

**Movie S1.** 1205Lu melanoma cells 1205Lu melanoma cells were transfected with: A483: aVBD; A524: aVBS1-2. Cells were perfused in a microfluidic chamber and allowed to adhere for 20 min prior to application of shear stress reaching 22.2 dynes.cm^-2^. The elapsed time is indicated in seconds.

